# The ratio between centromeric proteins CENP-A and CENP-C maintains homeostasis of human centromeres

**DOI:** 10.1101/604223

**Authors:** Daniël P. Melters, Tatini Rakshit, Minh Bui, Sergei A. Grigoryev, David Sturgill, Yamini Dalal

## Abstract

The centromere is the chromosomal locus that seeds the kinetochore, allowing for a physical connection between the chromosome and the mitotic spindle. At the heart of the centromere is the centromere-specific histone H3 variant CENP-A/CENH3. Throughout the cell cycle the constitutive centromere associated network is bound to CENP-A chromatin, but how this protein network modifies CENP-A nucleosome dynamics *in vivo* is unknown. Here, we purify kinetochore associated native centromeric chromatin and analyze its biochemical features using a combinatorial approach. We report that kinetochore bound chromatin has strongly reduced DNA accessibility and a distinct stabilized nucleosomal configuration. Disrupting the balance between CENP-A and CENP-C result in reduced centromeric occupancy of RNA polymerase 2 and impaired *de novo* CENP-A loading on the centromeric chromatin fiber, correlating with significant mitotic defects. CENP-A mutants that restore the ratio rescue the mitotic defects. These data support a model in which CENP-C bound centromeric nucleosomes behave as a barrier to the transcriptional machinery and suggest that maintaining the correct ratio between CENP-A and CENP-C levels is critical for centromere homeostasis.

## Introduction

The kinetochore is a large proteinaceous complex which physically connects centromeric chromatin to the mitotic microtubule spindles. Inaccuracies in kinetochore assembly can lead to the formation of dicentric chromosomes, or chromosomes lacking kinetochores. In either case, chromosomes fail to segregate faithfully, which drives genomic instability. Electron microscopy studies of mitotic centromeres reveal a two-layered electron dense structure that is over 200 nm in width and over 50 nm in depth (Comings & Okada, 1970; Rattner *et al*, 1975; Esponda, 1978; McEwen *et al*, 1998), delineated into the inner and outer kinetochore.

At the base of the inner kinetochore is the histone H3 variant CENP-A (CENH3), which is marked by its rapid evolution (Malik & Henikoff, 2001, 2003; Talbert *et al*, 2004; Cooper & Henikoff, 2004; Meraldi *et al*, 2006; Maheshwari *et al*, 2015), and its association with equally rapid evolving centromere DNA (Melters *et al*, 2013). This enhanced rate of evolution of sequences underlying the essential and conserved function is commonly referred to as the centromere paradox (Henikoff *et al*, 2001). Nevertheless, despite lack of sequence conservation at the level of CENP-A, and its associated DNA, in most species, CENP-A chromatin provides the epigenetic and structural foundation to assemble the kinetochore, recruiting inner kinetochore proteins CENP-B, CENP-C, and CENP-N (Régnier *et al*, 2005; Mendiburo *et al*, 2011). Together, these inner kinetochore components provide recognition motifs for outer kinetochore proteins in the conserved KMN network (KLN1, MIS12 and NDC80 complexes (Cheeseman *et al*, 2006; Przewloka *et al*, 2007; DeLuca & Musacchio, 2012; Weir *et al*, 2016). Deleting either CENP-A or CENP-C results in cell death or senescence (Fukagawa *et al*, 1999; Kwon *et al*, 2007; McKinley & Cheeseman, 2016), but this happens only after a few cell cycles. These data indicate that both CENP-A and CENP-C are often present in excess for what is required to form a functional kinetochore for one cell cycle. Furthermore, CENP-A and CENP-C are long lived proteins guaranteeing faithful chromosome segregation even after their respective genes have been deleted (Kalitsis *et al*, 1998; Howman *et al*, 2000; Suzuki *et al*, 2011; Bodor *et al*, 2013; Smoak *et al*, 2016).

Despite major advances made in understanding the hierarchy of the centromere and kinetochore proteins (McKinley & Cheeseman, 2016; Milks *et al*, 2009; Cheeseman, 2014; Klare *et al*, 2015), little is known about the physical features of the inner kinetochore bound to centromeric chromatin in multicellular eukaryotes. Individual mitotic kinetochores were isolated from budding yeast by using a FLAG-tagged outer kinetochore component Dsn1 as bait (Gonen *et al*, 2012). Point centromeres of budding yeast are unique because they display a one-to-one correspondence, in which a single CENP-A containing nucleosome (Bloom & Carbon, 1982; Saunders *et al*, 1988; Furuyama & Henikoff, 2009; Kingston *et al*, 2011; Krassovsky *et al*, 2012; Furuyama *et al*, 2013; Henikoff *et al*, 2014; Díaz-Ingelmo *et al*, 2015) binds to a single microtubule via the kinetochore. In contrast, the human centromeres are regional centromeres comprised of megabase-sized α-satellite arrays (Waye & Willard, 1989; Rudd *et al*, 2005). Recent advances in super resolution microscopy suggest that each human centromere harbors ∼400 CENP-A molecules (Bodor *et al*, 2014), which eventually associate with only ∼17 mitotic microtubule spindles (Suzuki *et al*, 2015). This inner kinetochore chromatin is thought to be folded into a boustrophedon (Ribeiro *et al*, 2010; Vargiu *et al*, 2017), in which the number of CENP-A nucleosomes present appears to exceed the number needed for centromere function. In recent work, we reported that CENP-A nucleosomes alone are twice as elastic as H3 nucleosomes, and that CENP-C fragments can suppress and rigidify CENP-A motions, correlating with increased compaction of the centromeric chromatin fiber *in vitro* and *in vivo*. (Melters *et al*, 2019) Here, we wished to provide deeper insights into kinetochore bound CENP-A chromatin in human cells.

Using nanoscale, immunofluorescence imaging, and biochemical approaches, we report that we were able to purify at least two classes of CENP-A chromatin populations in human cells. One class of CENP-A nucleosomes was weakly associated with inner kinetochore proteins and possesses diminutive dimensions; whereas a second class of CENP-A nucleosomes co-elute and co-purify with inner kinetochore proteins and display stabilized configurations. Interestingly, overexpression of CENP-C alters the balance of these two CENP-A classes, resulting in overall reduced RNA polymerase 2 occupancy, reduced *de novo* loading of CENP-A nucleosomes, and extensive mitotic defects. CENP-A mutants that either serve as a sink for excess CENP-C or which are unable to bind CENP-C rescue these mitotic defects. Altogether, these data support a working model in which the balance between CENP-A and kinetochore components is critical for centromere homeostasis.

## Results

### Two classes of centromeric chromatin can be purified from human centromeres

In previous work, we purified human CENP-A nucleosomes under a range of conditions, and analyzed them by various biochemical, EM and AFM approaches (Dimitriadis *et al*, 2010; Bui *et al*, 2012). We occasionally noticed a small fraction of macromolecular complexes within CENP-A native chromatin-immunoprecipitation (N-ChIP), that were large, compacted, and refractory to standard nucleosome analysis (Dimitriadis *et al*, 2010; Bui *et al*, 2012). We were curious to examine these larger complexes. Therefore, we developed a gentle, native, serial chromatin-immunoprecipitation assay to purify CENP-C bound chromatin from HeLa cells (Supplemental Figure S1). We mildly digested nuclear chromatin with micrococcal nuclease (MNase), and performed N-ChIP by first pulling-down CENP-C bound chromatin and subsequent pulling-down any remaining CENP-A chromatin using ACA serum (Earnshaw & Rothfield, 1985) (Figure 1A).

**Figure 1.**
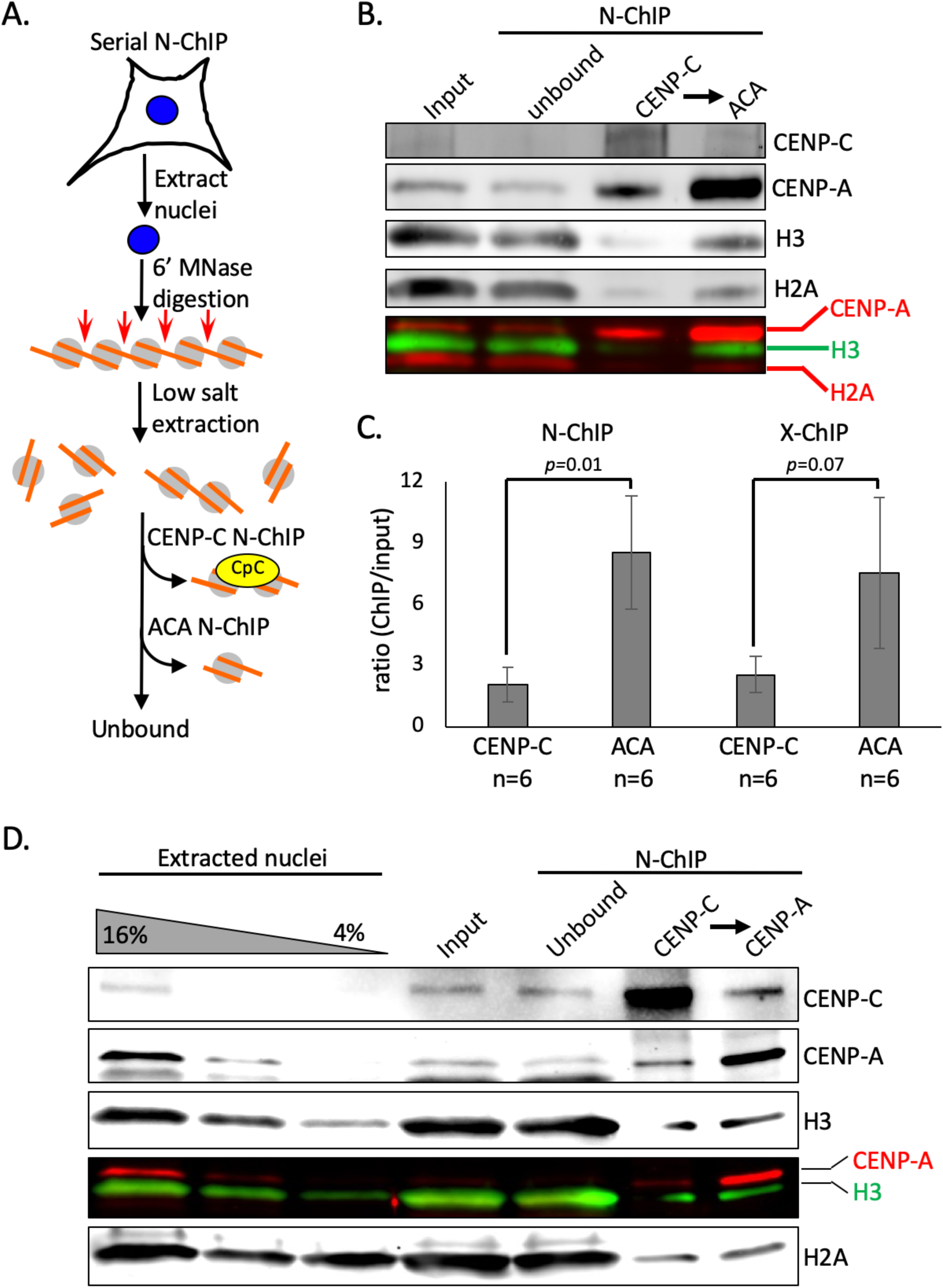
CENP-C binds a subset of CENP-A nucleosomes. (A) Schematic of experimental set-up of serial chromatin immunoprecipitation. (B) Western blot analysis of the serial N-ChIP was performed and probed for H2A, H3, CENP-A, and CENP-C. (C) Quantification of CENP-A enrichment in either CENP-C or subsequent ACA N-ChIP or X-ChIP confirms the presence of two CENP-A populations (paired t-test; significance was determined at *p*<0.05). (D) A serial dilution of extracted nuclei was loaded side-by-side to a CENP-C N-ChIP followed by a CENP-A N-ChIP.

We used quantitative immunoblotting to quantify the relative abundance of native CENP-A bound to CENP-C (Figure 1B) and free CENP-A chromatin. The enrichment of the N-ChIP’ed samples was measured over the input. We observed a 2.0±0.8-fold enrichment of CENP-A in the CENP-C N-ChIP and an 8.5±2.8-fold enrichment of CENP-A in the serial ACA N-ChIP (Figure 1C). Thus, our results indicate that in HeLa cells, there is a ∼4.7-fold excess of CENP-A that is weakly or not associated with CENP-C.

We wanted to test the possibility that during the experimental procedure of N-ChIP, weakly bound CENP-C could dissociate from CENP-A chromatin, we cross-linked samples prior to ChIP. Although in our hands overall chromatin extraction efficiency is generally lower under X-ChIP conditions, we still observed a 2.5±0.9-fold enrichment of CENP-A in the CENP-C X-ChIP and a 7.5±3.7-fold enrichment of CENP-A in the ACA X-ChIP (Figure 1C, Supplemental Figure S1E, Table S1).

We next tested the possibility that CENP-C may be dissociating from one class of CENP-A chromatin, in which case it would accumulate in the nuclear pellet. To examine how much CENP-C is lost during purification, we loaded a gradient of nuclear extract in addition to the serial N-ChIP (Figure 1D). We observed that very little CENP-C remained in the nuclear debris/pellet. These data lend confidence to the interpretation that the majority of CENP-C was pulled-down in the CENP-C N-ChIP.

Next, we tested whether the two types of CENP-A populations have similar sedimentation patterns. Following MNase digestion, we ran the chromatin on a 5-20% glycerol gradient and performed serial N-ChIP on both every fraction (Supplemental Figure S2A). Whereas the free CENP-A population had a sedimentation pattern very similar to input chromatin, CENP-C associated CENP-A chromatin showed a distinctively different sedimentation pattern (Supplemental Figure S2). The relative abundance of longer chromatin arrays in the CENP-C N-ChIP was also observed by high resolution capillary electrophoresis (BioAnalyzer, Supplemental Figure S3). We were curious to test whether CENP-C-associated chromatin occupied unique centromeric sequences compared to CENP-C-depleted CENP-A chromatin. In order to test this possibility, we performed ChIP-seq replicates, and analyzed the DNA sequences against the several centromeric references. These analyses showed that both CENP-A populations occupy similar centromeric sequences (Supplemental Figure S4).

Altogether, these data suggest that there is a sizeable fraction of CENP-A chromatin which is weakly associated with CENP-C, and a smaller fraction of CENP-A which is very robustly bound to CENP-C.

### CENP-A nucleosomes robustly bound to CENP-C co-purify with other CCAN components

The chromatin we extracted is from cycling HeLa cells, the majority of which are in G1 phase (Pettersen *et al*, 1977). Throughout the cell cycle the constitutive centromere associated network (CCAN) complex, which is composed of several proteins, is bound to the centromeric chromatin to form the inner kinetochore (Figure 2A) (Pesenti *et al*, 2016; McKinley & Cheeseman, 2016). To test whether other components of the CCAN are present within the complex within CENP-A:CENP-C above, we performed western blot analyses on these purified CENP-C complexes. In addition to CENP-A, H2A, and H2B, CCAN components CENP-B, CENP-I, CENP-N, CENP-W, and CENP-T were enriched (Figure 2B). To our surprise, the dedicated chaperone for CENP-A assembly, HJURP was also enriched in the CENP-C N-ChIP. This result supports a recent report from novel BioID experiments, which show that HJURP is associated with CENP-A chromatin at multiple points of the cell cycle (Remnant *et al*, 2019). Overall, our purified CENP-A:CENP-C complex robustly represents the CCAN. Furthermore, outer kinetochore components MIS12 and HEC1/NDC80 were modestly enriched in the CENP-C N-ChIP (Figure 2B). Altogether, from these data, we deduced that the large CENP-C complex that was enriched in fraction 12 of our glycerol gradient experiments (Supplemental Figure S2D) includes the full CCAN, possibly representing fully matured kinetochores.

**Figure 2.**
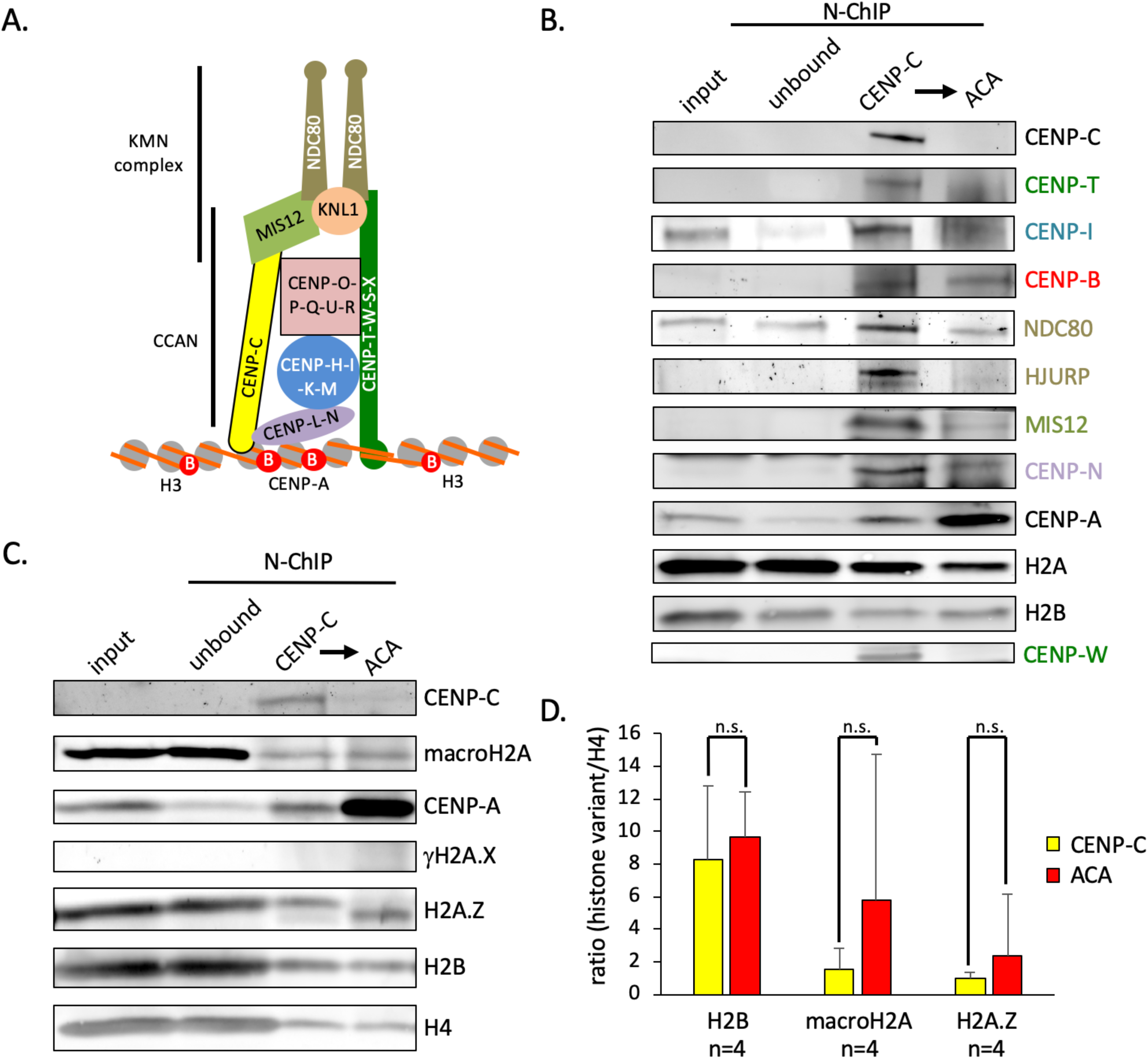
A small fraction of CENP-A stably associates with the inner kinetochore. (A) To determine if CENP-C N-ChIP also brought down kinetochore components, both CENP-A populations were purified and probed for the histones (H2A, H4, CENP-A), CENP-B, inner kinetochore components (CENP-W, CENP-T, CENP-I, CENP-N), and outer kinetochore components (MIS12, HEC1). (C) To determine whether histone H2A variants associate with CENP-A nucleosomes in complex with CENP-C or bulk CENP-A, we performed western blot analysis probing for γH2A.X, H2A.Z, and macroH2A. (D) Quantification of four independent experiments revealed that none of the H2A variants was enriched in either CENP-A population. As a control, we measured the relative amount of H2B over H4 (paired t-test; significance was determined at *p*<0.05).

### Inner kinetochore associated chromatin is not uniquely enriched in histone H2A variants

Heterotypic CENP-A/H3.3 nucleosomes have been reported, but they are generally restricted to ectopic sites (Athwal *et al*, 2015; Nye *et al*, 2018; Lacoste *et al*, 2014). Where histone H3 and H2B have few variants, there are several histone H2A variants including macroH2A, H2A.Z, and γH2A.X (Melters *et al*, 2015). Indeed, mass-spectrometry data has suggested that H2A histone variants might form nucleosomes with CENP-A (Foltz *et al*, 2006; Bailey *et al*, 2016). We therefore set out to interrogate the possibility that CENP-A nucleosomes associated with the inner kinetochore contained other histone variants. As before, we performed a CENP-C N-ChIP followed by a serial ACA N-ChIP and tested both samples for the presence of histone H2A variants. Interestingly, while we detected H2A.Z and macroH2A variants in both CENP-C and ACA N-ChIP (Figure 2C), their relative abundance was not quantifiably different between the two CENP-A populations (Figure 2D, Supplemental Table S2). Consequently, while these data do not currently yield evidence that H2A variants might contribute to structural differences between the two CENP-A populations, they do support previous data (Foltz et al,. 2006; Bailey et al., 2016), suggesting that H2A variants can incorporate into centromeric chromatin, albeit by mechanisms yet unknown.

### Development of immuno-AFM to verify the identity of nucleosomes associated with the CENP-C complex

The purification of CENP-A chromatin bound to the full CCAN from human cells presents an exciting opportunity to study this complex in its native form. In order to do this robustly, we needed a method to confirm the identity of nucleosomes associated with CENP-C, complementary to the Western blot analyses provided above (Figures 1, S2). To this end, we developed a single-molecule based method to test whether CENP-A nucleosomes are physically present in CENP-C N-ChIP samples. Inspired by classical immuno-EM protocols (e.g. Dimitriadis *et al*, 2010), and recognition-AFM (e.g. Wang *et al*., 2008), in which we have shown we can confirm the identity of histones in a purified biological sample, we adapted immuno-labeling for in-air atomic force microscopy (AFM) (Figure 3A).

**Figure 3.**
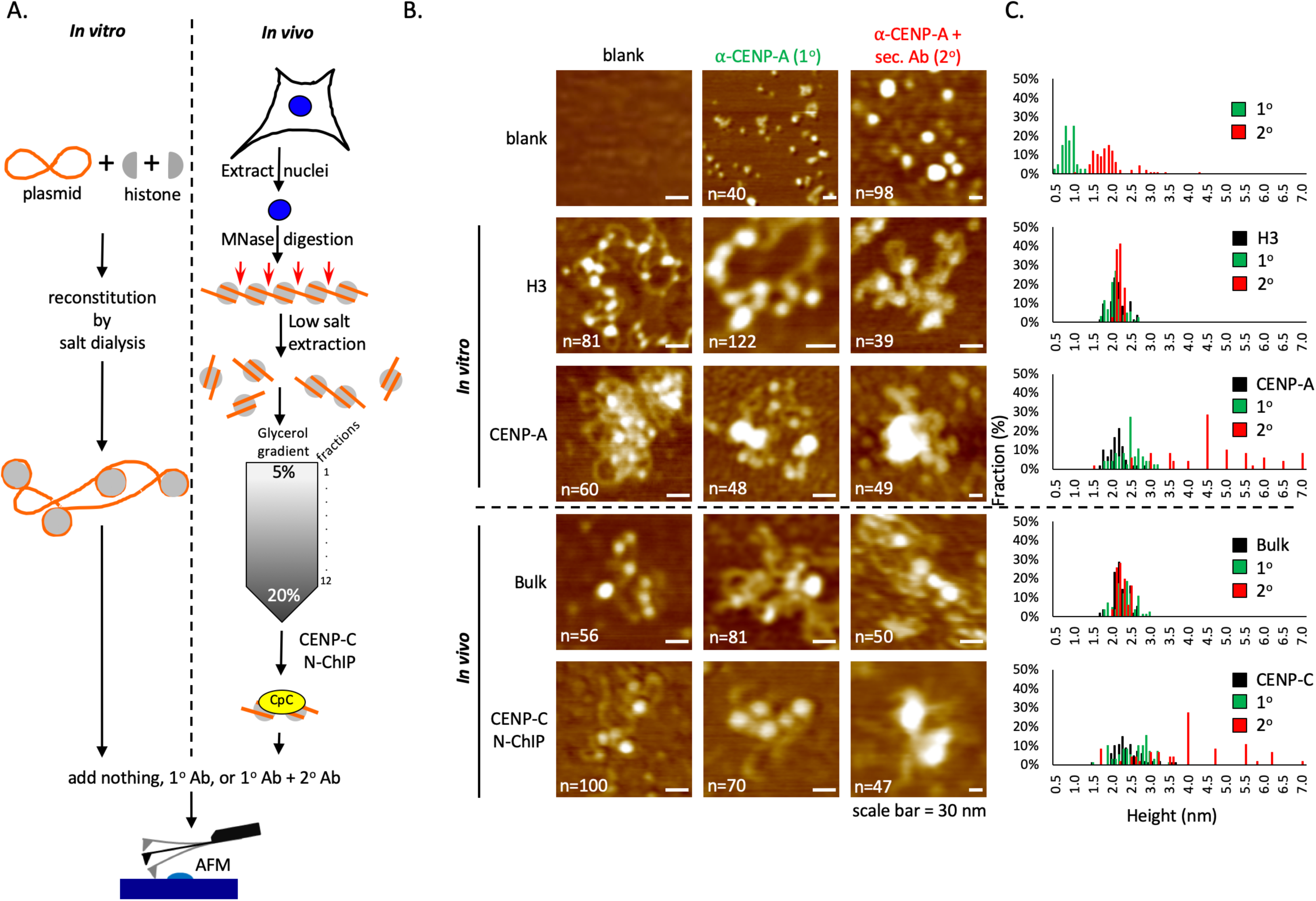
Immuno-AFM confirms CENP-C associates with CENP-A nucleosomes. (A) To confirm that CENP-C associated with CENP-A nucleosomes, we performed immuno-AFM on *in vitro* reconstituting H3 and CENP-A nucleosomes in parallel to extracted bulk and CENP-C associated chromatin. (B) Representative images for three conditions (no antibody, 1° antibody, 1° plus 2° antibodies) per sample. The scale bar is 30 nm. (C) Height measurement of all three conditions were plotted per sample, showing that anti-CENP-A antibody only recognized *in vitro* reconstituted CENP-A nucleosomes and nucleosomes associated with CENP-C, confirming that CENP-C indeed strongly associated with CENP-A nucleosomes.

We first visualized samples by AFM either without antibody; or, primary (1°) mouse monoclonal anti-CENP-A antibody; or, 1° CENP-A antibody and, either secondary (2°) anti-mouse antibody or 2° anti-mouse Fab fragment. The 1° antibody by itself was 0.8±0.2 nm in height and the addition of the 2° antibody resulted in a height increase to 2.0±0.5 nm (Figures 3B, C, Supplemental Table S3, Raw Data File 1). To confirm the 1° antibody’s specificity, we used *in vitro* reconstituted recombinant H3 or CENP-A nucleosomes as before and incubated them with either no antibody (no Ab); 1° alone, or 1° + 2° antibodies, respectively (Figures 3B, C). As expected, control *in vitro* reconstituted H3 nucleosomes did not show a shift in particle height in the presence of anti-CENP-A antibodies (no Ab: 2.2±0.2nm, 1°: 2.1±0.2 nm, and 2°: 2.2±0.1 nm, resp.). However, *in vitro* reconstituted CENP-A nucleosomes did increase in height upon binding to their antibodies (no Ab: 2.2±0.2 nm, 1°: 2.5±0.3 nm, and 4.6±1.4 nm with 2° antibody, or 3.2±0.6 nm with Fab fragment, resp., Figures 3B, C, S5, Supplemental Table S3, Raw Data File 1).

Having standardized this approach on *in vitro* reconstituted samples, we next applied this method to *in vivo* samples, namely native H3 or CENP-C purified chromatin (Figures 3B, C). Similar to reconstituted H3 nucleosomes, native bulk H3 chromatin did not demonstrate a shift in particle height when incubated with anti-CENP-A antibodies (no Ab: 2.3±0.2 nm, 1°: 2.4±0.3 nm, and 2°: 2.3±0.1 nm, resp.). In contrast, nucleosomal particles that came down in the CENP-C N-ChIP displayed a shift in height when challenged with anti-CENP-A antibodies (no Ab: 2.4±0.4 nm, 1°: 2.6±0.4 nm, and 2°: 3.9±1.3 nm, resp.). These results support biochemical evidence based on Western blots, that CENP-A nucleosomes are indeed associated with the purified CENP-C complex in these purified complexes.

### Physically altered CENP-A nucleosomes are bound to the inner kinetochore

We next sought to analyze physical characteristics of purified CCAN associated CENP-A chromatin in its native context. This, to our knowledge, has not been accomplished before from native human centromeres. We split the samples in half and imaged the same samples independently (by two different operators) using two complementary high resolution single-molecule methods: in-air AFM, and transmission electron microscopy (TEM) (Figures 4A, B, S6). To retain integrity of the chromatin, samples were analyzed within 24 hours of purification. By AFM and by TEM, we observed several similar features. First, we observe large polygonal structures with a roughly circular footprint (height: 5.8±2.1 nm; area: 629±358 nm^2^), generally associated with four to six nucleosomes (Figures 4A, B, S6, Table S4, Raw Data File 2). Compared to individual CENP-A nucleosomes, these complexes were substantially larger. As important controls, we analyzed control mock IPs, bulk chromatin, or CENP-A N-ChIP (Figures 4A, B, S6), which did not display these polygonal complexes. Together, this suggests these structures arise specifically from the CENP-C/CCAN bound complex. Interestingly, by AFM, we observed that the largest of CENP-C complexes tends to be associated with a ∼230bp long nucleosome-free DNA (Figure 4A, Table S4, Raw Data File 2). The same samples, analyzed in parallel by TEM, displayed similar features (Figure 4B), including the association of four to six nucleosomes at the periphery of the complex.

**Figure 4.**
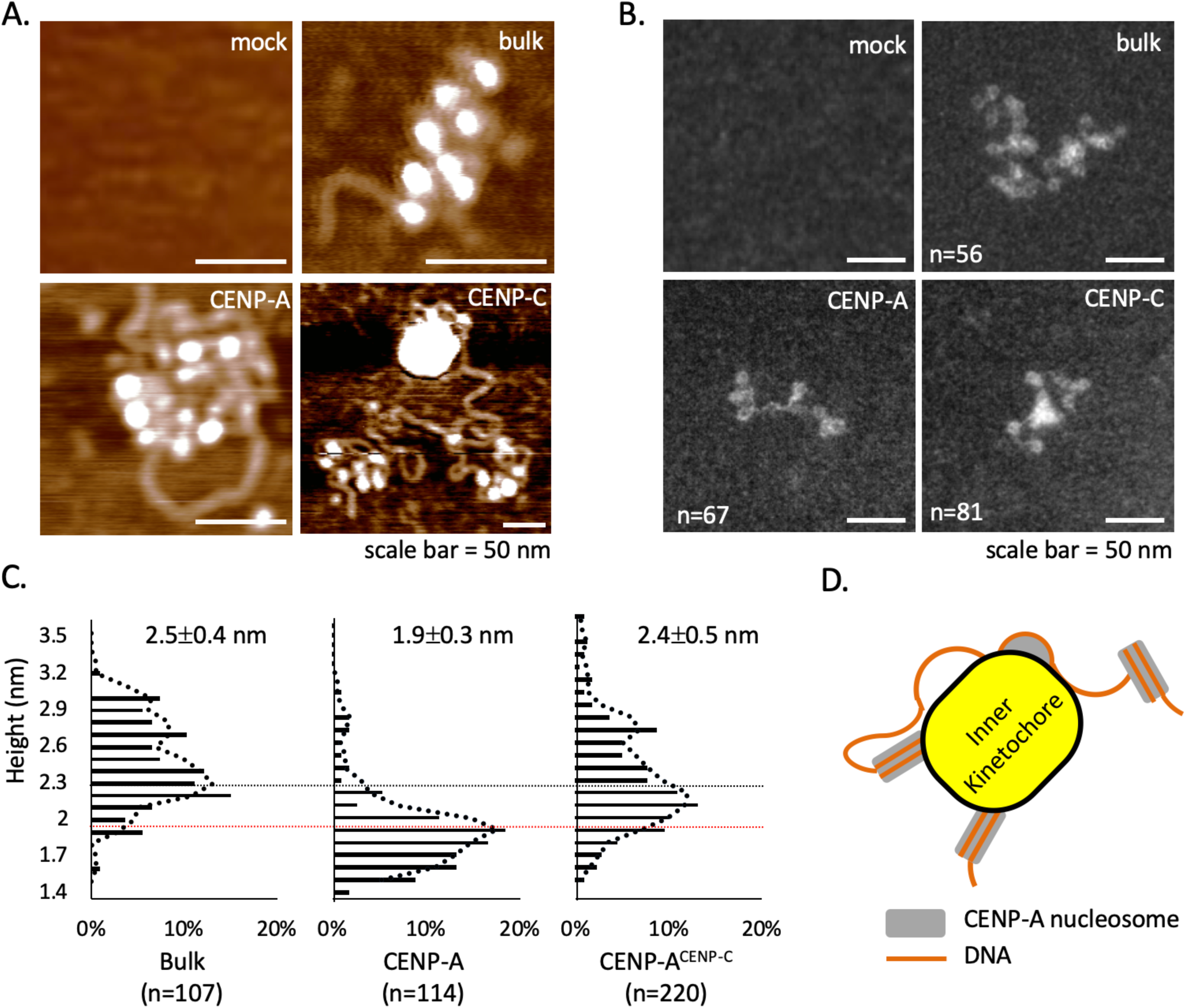
Purified CENP-C complex are associated with stable octameric CENP-A nucleosomes. Chromatin was extracted from HeLa cells after 6-minute MNase digestion, followed by either mock, CENP-A, or CENP-C N-ChIP. Unbound chromatin was used for bulk chromatin. Representative AFM (A) and TEM (B) images of either mock, bulk chromatin, CENP-A chromatin, or CENP-C chromatin showed that CENP-C complex is a large polygonal structure. (C) Nucleosomal height was quantified for either bulk chromatin, CENP-A, or CENP-A nucleosomes associated with CENP-C complex. The mean height with standard error is shown, as well as their distribution. Robustly kinetochore associated CENP-A nucleosomes were significantly taller than weakly or not bound CENP-A nucleosomes (2-sided t test *p*<2.6×10^-18^). (D) A model showing how 4-6 CENP-A nucleosomes associate with the inner kinetochore, including a stretch of 230-bp of naked DNA that is refractory to MNase digestion.

We next turned our attention to individual nucleosomes associated with the CENP-C complex. In our previous work, *in vitro* reconstituted recombinant octameric CENP-A nucleosomes on 601 or α-satellite DNA behave similar to H3 when measured by in-air AFM, with average heights of ∼2.4 nm and widths of ∼12 nm (Athwal *et al*, 2015; Walkiewicz *et al*, 2014b). This observation is consistent with measurements made by static biophysical, EM, and crystallographic methods, which generally show that except for flexible exit/entry DNA termini and loop 1, *in vitro* reconstituted CENP-A nucleosomes have dimensions similar to H3 nucleosomes (Tachiwana *et al*, 2011; Walkiewicz *et al*, 2014b; Roulland *et al*, 2016; Kim *et al*, 2016; Vlijm *et al*, 2017).

In contrast, native CENP-A nucleosomes purified from either fruit fly or human cell lines generally display smaller dimensions compared to native H3 nucleosomes, except during S phase (Athwal *et al*, 2015; Bui *et al*, 2012; Dimitriadis *et al*, 2010; Dalal *et al*, 2007; Bui *et al*, 2013). Indeed, we recapitulated these observations here. Relative to H3 nucleosomes (2.5±0.3 nm) (Figure 4C, Table S5, Raw Data File 3), native bulk CENP-A nucleosomes not associated with CENP-C are continuing to be uniquely identifiable by their smaller average height of 1.9±0.3 nm (Figure 4C).

We examined CENP-A nucleosomes that are physically associated with the CENP-C complexes (Figure 4C, Table S5, Raw Data File 3). CENP-A nucleosomes associated with CENP-C complexes had distinctly larger dimensions, with a height of 2.4±0.5 nm (Figure 4C), significantly taller than bulk CENP-A nucleosomes alone (two-sided t-test p < 0.001).

These results point to the presence of two physically distinct CENP-A nucleosomes within the human centromere: one species of CENP-A nucleosome which is shorter, and another species of CENP-A nucleosome associated with the CENP-C/CCAN complex (Figure 4D), which adopts a taller configuration.

### CENP-C overexpression does not lead to increased DNA breaks

Cumulatively, the data above suggest that one type of structure in the inner kinetochore is comprised of a polygonal dome-like structure with a roughly circular footprint, associated with a chromatin sub-domain comprised of four to six octameric CENP-A nucleosomes. Yet another type of domain in the centromere contains smaller CENP-A nucleosomes. Chromatin fiber studies show that CENP-A and H3 chromatin domains are interspersed (Sullivan & Karpen, 2004; Blower *et al*, 2002; Ribeiro *et al*, 2010; Kyriacou & Heun, 2018). Indeed, recent work show that CENP-C and CENP-A do not perfectly overlap on chromatin fibers (Padeganeh *et al*, 2013; Kyriacou & Heun, 2018). Recent structural studies of CENP-A nucleosome suggest that CENP-A nucleosome are innately more flexible compared to H3 nucleosomes (Winogradoff *et al*., 2015; Falk *et al*., 2016; Roulland *et al*., 2016; Malik *et al*., 2018; Melters *et al*., 2019 in press), whereas CENP-C strongly suppresses CENP-A nucleosomes motions (Falk *et al*., 2015, 2016; Melters *et al*., 2019 in press). This raises an intriguing question on potential roles for maintaining unbound or free CENP-A centromeric chromatin *in vivo*.

A plausible and attractive hypothesis is that during mitosis, flexible CENP-A particles might provide a mechanical “bungee”-like state, which allows the dissipation of mitotic forces. In this scenario, excess CENP-C would be predicted to dampen the motions of CENP-A nucleosomes, thereby reducing the overall springiness of centromeric chromatin. Thus, loss of the flexible CENP-A domain might result in an accumulation of DNA breaks during mitosis. To test this hypothesis, we overexpressed C-terminally tagged GFP CENP-C (CENP-C^OE^) for three days, in cells synchronized to late mitosis and early G1 and scored for the DNA break marker γH2A.X. Although we found an increase in mitotic defects, no appreciable increase in γH2A.X foci was observed at centromeric foci (Figure 5). These data suggest that partial reduction in the amount of free CENP-A does correlate with mitotic defects, a priori it does not appear to cause an increase in DNA breaks. We were curious why these mitotic defects arose and considered alternative hypotheses that might explain the result.

**Figure 5.**
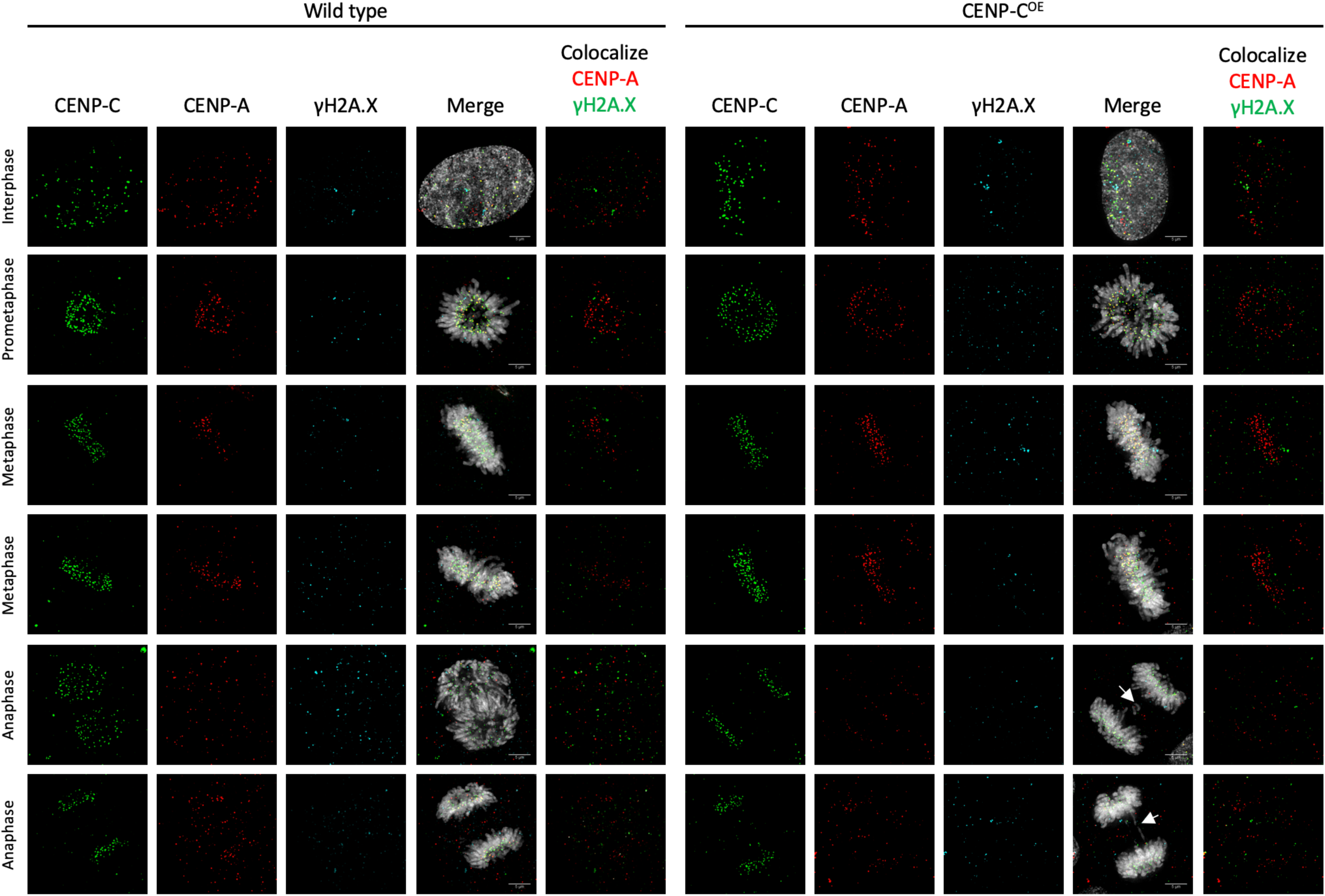
CENP-C overexpression did not increase mitotic double strand DNA breaks. Representative images of either wild-type or CENP-C overexpressing HeLa cells. HeLa cells were transfected for three days and synchronize to early G1 prior to fixation and staining for CENP-C (green), CENP-A (red), γH2A.X (cyan), and DAPI (grey). No difference in number of γH2A.X foci was observed between the two samples. Arrow highlight lagging chromosomes.

### CENP-C overexpression limits *de novo* CENP-A loading

An obvious second hypothesis arises from decades of data in the transcription field, which suggests that chromatin that is more “open” or accessible, is more likely to be transcriptionally permissive (Maeshima *et al*, 2019; Klemm *et al*, 2019). Thus, we speculated that free CENP-A chromatin might be necessary for centromeric transcription. RNA polymerase 2 (RNAP2) mediated centromeric transcription has been shown to be critical for *de novo* CENP-A loading in multiple species (reviewed in Müller and Almouzni, 2017). In parallel work, we have observed that overexpression of CENP-C results in reduced RNAP2 levels at centromeric chromatin (Melters *et al*., 2019 in press). Therefore, a logical prediction is that limiting access of the transcriptional machinery to CENP-A chromatin (Figures 6A, B), should reduce new CENP-A loading. An initial clue supporting this possibility was deduced from western blot analysis, in which over-expression of CENP-C led to a significant reduction in the free CENP-A population (two-sided t-test *p*<0.05; Figure 6B, Table S6). To test whether CENP-C^OE^ would specifically lead to a reduction in new CENP-A loading, we turned to the well-established SNAP-tagged CENP-A system combined with quench pulse-chase immunofluorescence (Bodor *et al*, 2012). Using this system in cells synchronized to mid-G1, one can distinguish between older CENP-A (TMR-block) and newly incorporated CENP-A (TMR-Star) (Figures 6C, D). Strikingly, in the CENP-C overexpression background, in which we observed RNAP2 is depleted from centromeric chromatin in early G1 (Figure 6A, B), we concomitantly observed a 2.3-fold reduction of *de novo* incorporation of CENP-A (two-sided t-test *p*<0.00001; Figures 6E, F, Raw Data File 4). Therefore, *in vivo* CENP-C^OE^ leads to suppression of RNAP2 occupancy, reduction in total free CENP-A levels (Figures 6A, B), and a reduction in *de novo* CENP-A loading (Figures 6E, F).

**Figure 6.**
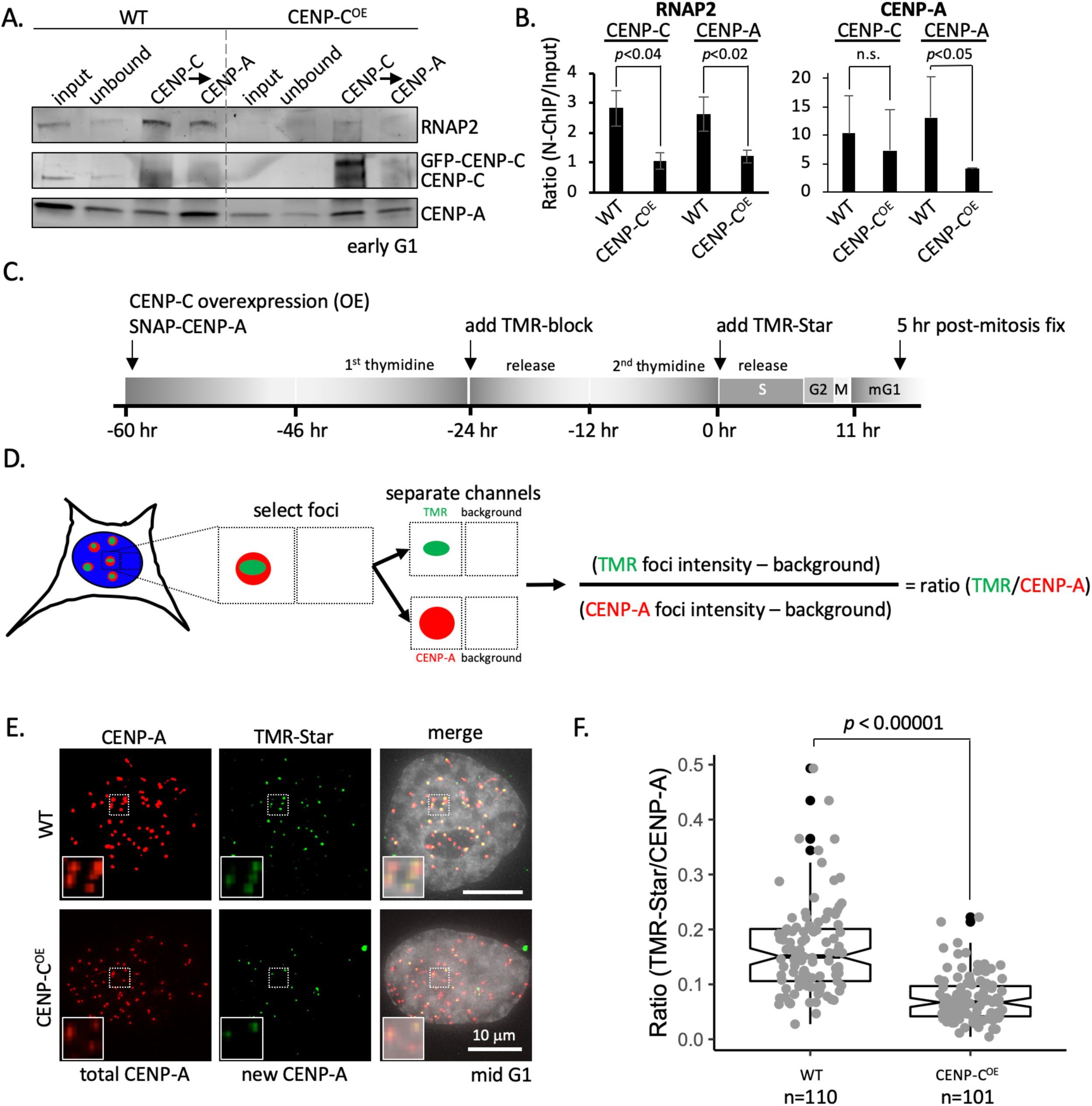
New CENP-A loading impaired upon CENP-C overexpression. (A) Western blot comparing wild-type versus CENP-C^OE^ conditions for RNAP2, CENP-C, and CENP-A levels at early G1. (B) Quantification of RNAP2 and CENP-A levels. (C) Schematic of experimental design. (D) Colocalizing immunofluorescent signal for CENP-A and TMR-Star are collected and the intensity of both foci is measured as well as background directly neighboring the foci to determine the ratio TMR-star signal over total CENP-A signal. (E) *De novo* CENP-A incorporation was assessed by quench pulse-chase immunofluorescence. After old CENP-A was quenched with TMR-block, newly loaded CENP-A was stained with TMR-Star and foci intensity was measured over total CENP-A foci intensity. Inset is a 2x magnification of the dotted box in each respective image. (F) Quantification of *de novo* CENP-A loading by measuring the ratio of TMR-Star signal over total CENP-A signal.

### CENP-A mutants rescue mitotic defects caused by CENP-C overexpression

As has been reported previously in chicken DT-40 cells (Fukagawa *et al*, 1999), we observed that overexpressing CENP-C resulted in a quantifiable increase in mitotic defects (40% normal, 60% abnormal) relative to wildtype cells (74% normal, 36% abnormal), most notably lagging chromosomes and multipolar spindles (Figure 7A, B).

**Figure 7.**
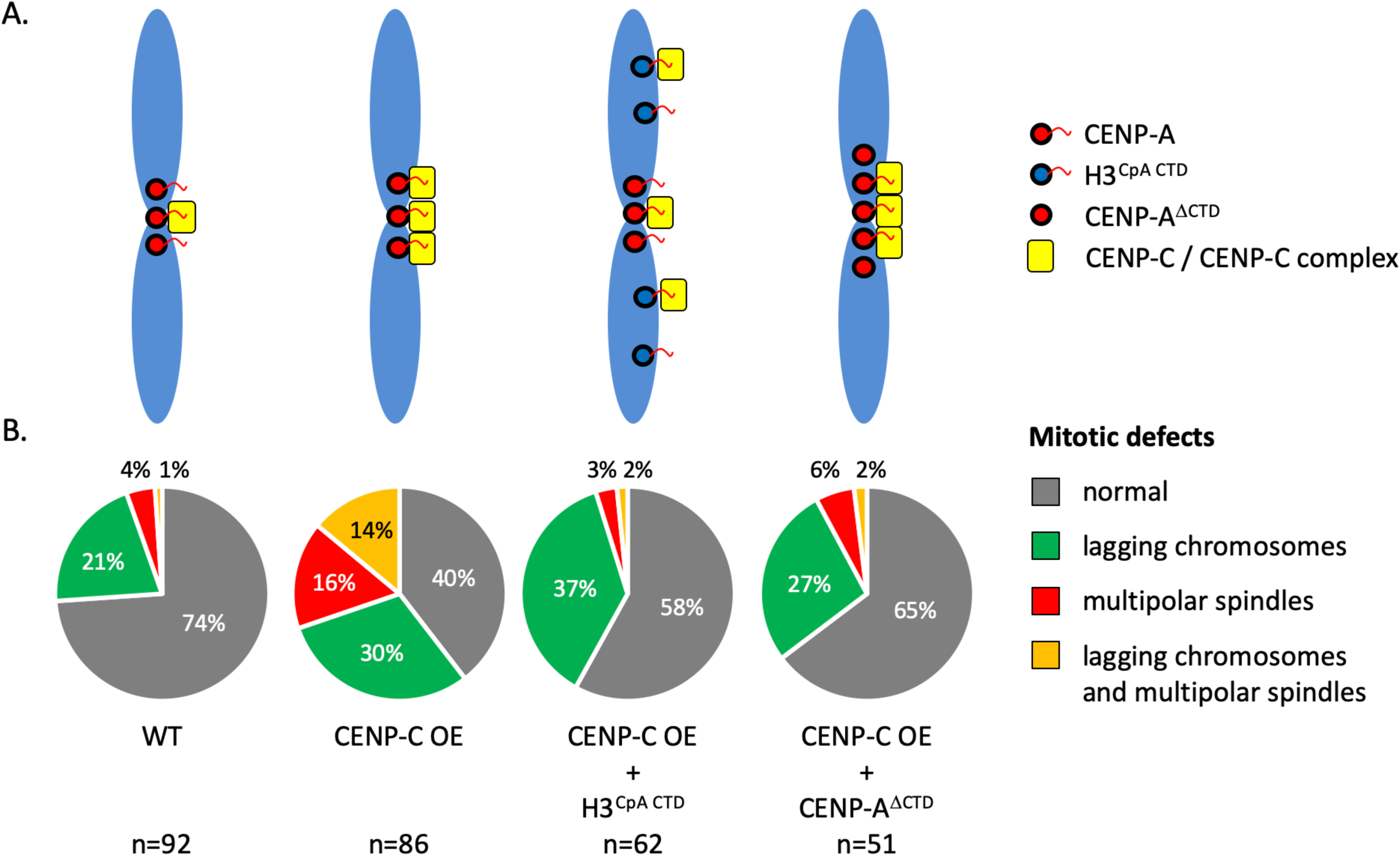
CENP-C overexpression leads to increased mitotic defects, which can be rescued with CENP-A CTD mutants. (A) Three days of ectopic overexpression of CENP-C, which were synchronized to M phase, resulted in dramatic increase in mitotic defects compared to wild-type cells. The C-terminal tail of CENP-A is essential for recruitment of CENP-C. We reasoned that to rescue the mitotic defect of CENP-C overexpression by co-expressing histone H3 with the C-terminal tail of CENP-A (H3^CpA CTD^) or CENP-A lacking its C-terminal tail (CENP-A^ΔCTD^). (B) Mitotic defects were quantified. We observed that the level of both multipolar spindle (red) and multipolar spindle with lagging chromosome (orange) were reduced to wild-type levels.

CENP-C functionally docks at the C-terminal tail of CENP-A nucleosomes (Carroll *et al*, 2010; Falk *et al*, 2016, 2015; Kato *et al*, 2013; Guo *et al*, 2017; Ali-Ahmad *et al*, 2019). We reasoned that expressing CENP-A mutants which can either sequester away excess CENP-C, or which are insensitive to CENP-C, should rescue the defects noted above. Therefore, in the background of CENP-C over-expression, we expressed either a fusion of H3 with the C-terminal tail of CENP-A (H3^CpA CTD^) which can bind CENP-C, or CENP-A lacking its C-terminal tail (CENP-A^ΔCTD^) which can still be deposited to centromeres by the chaperone HJURP, but which cannot bind CENP-C (Figure 7A).

In the background of CENP-C overexpression, H3^CpA CTD^ should function as a sink for excess CENP-C. In contrast, CENP-A^ΔCTD^ should reintroduce a free CENP-A population. Indeed, upon scoring mitotic cells in the background of overexpressing CENP-C and the two mutants, we observed that multipolar spindle defects were rescued by both H3^CpA CTD^ (58% normal, 42% abnormal), and CENP-A^ΔCTD^ (65% normal, 35% abnormal) (Figure 7B).

Together, these data show that CENP-C overexpression reduces RNAP2 occupancy at the centromeric chromatin, reduces free CENP-A levels (Figures 6A, B), and reduces incorporation of *de novo* CENP-A (Figure 6E, F), and results in an increase in mitotic defects (Figure 7B). Reintroducing either a free CENP-A population or a H3^CpA CTD^ sink results in a rescue of mitotic defects (Figure 7B). These data suggest a working model where a balance between kinetochore bound and free CENP-A chromatin is important for centromere homeostasis.

## Discussion

Since the discovery of CENP-A (Earnshaw & Rothfield, 1985; Palmer *et al*, 1991), it has been demonstrated that CENP-A nucleosomes are required and sufficient to form kinetochores (Régnier *et al*, 2005; Mendiburo *et al*, 2011). At the same time, it is puzzling that more CENP-A nucleosomes reside at the centromere than are strictly needed to successfully seed a kinetochore (McKinley & Cheeseman, 2016; Bodor *et al*, 2013). Indeed, while our current study was under revision, a recent chromatin fiber study employed a proximity based labeling technique called BioID, finding that CENP-A foci do not entirely overlap with CENP-C foci (Kyriacou & Heun, 2018). Our data strongly supports and extends this finding using a completely independent approach.

We report on the existence of two classes CENP-A nucleosomes *in vivo*. One class of CENP-A nucleosomes was strongly associated with the CENP-C complex, along with other components of the inner kinetochore/CCAN; whereas another population of CENP-A nucleosomes is weakly associated with CENP-C. Strikingly CENP-A nucleosomes associated with the CENP-C complex have altered dimensions relative to bulk CENP-A nucleosomes alone (Figure 4D). All-atom computational modeling has previously indicated that CENP-A nucleosomes have a weaker four helix bundle resulting in an intrinsically more disordered nucleosome core compared to a canonical H3 nucleosome (Winogradoff *et al*, 2015), that this flexibility arises from the CENP-A/H4 dimer (Zhao *et al*, 2016), is lost in the tetramer (Zhao *et al*, 2019) but regained in the CENP-A octamer, resulting in an energetically “frustrated” nucleosome, which predicts the existence of multiple deformation states of CENP-A (Winogradoff *et al*., 2015; Bui *et al.,* 2012). These data also predict that kinetochore components especially CENP-C could fix one preferred conformational state. Indeed, previous *in vitro* work has shown that both the central domain and the conserved CENP-C motif make contact with the CENP-A C-terminal tail and H2A acidic patch (Kato *et al*, 2013). In hydrogen-deuterium exchange and smFRET experiments of *in vitro* reconstituted CENP-A mononucleosomes, these contacts result in changes in the overall shape of CENP-A nucleosome by bringing the two nucleosome halves closer together and limiting the DNA sliding (Falk *et al*, 2015, 2016; Guo *et al*, 2017). As a result, the CENP-A nucleosome when bound to the central domain of CENP-C has an appearance similar to a canonical H3 nucleosome, a result we recently demonstrated using computational, nanobiophysical and biochemical tools (Melters *et al*, 2019). Indeed, in the CENP-C purified samples from human centromeres, we observe a shift in nucleosomal height when CENP-A nucleosomes associate with CENP-C complexes. One interpretation of this shift is that the CENP-C complex stabilizes the octameric conformation of the CENP-A nucleosome *in vivo*. We note that other interpretations are equally plausible: for example, the binding of kinetochore partners may alter the physical dimensions of nucleosomes in the complex; or, the absence or presence of DNA/histone modifications might limit the conformational space that can be sampled on minute time scales by CENP-A nucleosomes (Winogradoff *et al*, 2015; Bui *et al*, 2017; Stumme-Diers *et al*, 2018).

Interestingly, overexpression of CENP-C in HeLa cells induced a 6-fold increase in multipolar spindles and 2-fold increase in lagging chromosome (Figure 7). A correlation between multipolar spindles and merotelic chromosome attachment was described (Silkworth *et al*, 2009). Merotelic attachment are a common feature of cancer cells and are believed to be a driving force of chromosome instability (CIN) (Salmon *et al*, 2005; Gregan *et al*, 2011). Interestingly, both the multipolar and the lagging chromosome phenotype we observed were rescued by re-establishing a balance between the two CENP-A domains (Figure 7). This raises the possibility that multipolar spindles can be directly regulated by either the kinetochore or possibly even CENP-C. Several important questions arise from this observation; first, does overexpression of CENP-C create a platform which might allow a larger kinetochore to be formed, thereby promoting merotelic attachments; second, does free CENP-A chromatin provide cues to direct the kinetochore to face the spindle poles, either by organizing centromeric chromatin and/or by regulating inner centromere proteins localization, like Aurora B? These will be interesting questions to pursue in follow-up work.

We observed that overexpression of CENP-C also reduced the level of *de novo* incorporation of CENP-A nucleosomes (Figure 6E, F). Previous work has shown that CENP-A incorporation is dependent on centromeric transcription (reviewed in Müller and Almouzni, 2017). Indeed, levels of centromeric RNAP2 were reduced in early G1 (Figure 6B), providing further evidence that nucleosome loading requires polymerase activity. Furthermore, recent work showed that nucleosome binding partners have the capacity to alter nucleosome’s innate distortability (Falk *et al*, 2016, 2015; Sanulli *et al*, 2019), as well as compact chromatin (Melters *et al*, 2019; Sanulli *et al*, 2019). Altogether, this points to the intriguing possibility that CENP-C overexpression alters centromeric chromatin is such a way that it becomes less accessible to the transcriptional machinery. This fraction of chromatin is in juxtaposition to free CENP-A chromatin, which is unable to be bound and suppressed by linker histone H1 (Roulland *et al*, 2016), combined with the intrinsic distortability of CENP-A nucleosomes might (Winogradoff *et al*., 2015; Malik *et al*., 2018), we think, create an innate open chromatin state, readily accessible to specific transcription associated factors. Nevertheless, there are some caveats which present a puzzle for which we have not arrived at satisfying answers yet. In wild-type conditions we observed RNAP2 to be present in both CENP-A populations. This raises the question, if CENP-C associated CENP-A chromatin is less accessible, how did RNAP2 get there? One feasible answer is that the entry and exit DNA strands of CENP-C associated CENP-A nucleosomes are as accessible as in unbound CENP-A nucleosomes; but access to the DNA wrapped by the CENP-A nucleosome is impaired upon CENP-C binding. To load RNAP2 to the chromatin fiber, the pre-initiation complex (PIC) has to be formed first. Thus, an important current problem is to identify where the PIC forms-at the free CENP-A chromatin, or on the bordering H3K4me2 containing domains mapped within the centromere over two decades ago, the function of which still remains mysterious (Sullivan & Karpen, 2004).

In summary, in this report we describe that CENP-A nucleosomes display two alternative conformational states *in vivo,* both of which are important for centromere fidelity in cycling cells. Nucleosome dynamics play an important role in genome compaction, protection from DNA damaging agents, and regulating DNA access by DNA binding factors. These dynamics are driven by only a few interactions between the interfaces of DNA and nucleosomes (Polach & Widom, 1995; Widom, 1998; Fierz & Poirier, 2019). Recently, we described a CENP-A core post-translation modification (PTM) that altered the binding of CENP-C to CENP-A *in vivo* (Bui *et al*, 2017). An exciting line of future investigation is to examine how DNA or histone modifications which promote specific conformations of CENP-A nucleosomes, might alter its interactions with chaperones, kinetochore partners, and its occupancy on centromere α-satellite DNA. It will also be of interest to investigate whether nucleosome stabilizing or destabilizing interactions promote or suppress centromeric transcription required for the epigenetic memory of centromeres in various species (Wong *et al*, 2007; Chueh *et al*, 2009; Li *et al*, 2008; Ferri *et al*, 2009; Bergmann *et al*, 2011; Choi *et al*, 2011; Ohkuni & Kitagawa, 2011; Quénet & Dalal, 2014; Rošić *et al*, 2014; Catania *et al*, 2015; Bobkov *et al*, 2018; McNulty *et al*, 2017; Zhu *et al*, 2018; Ling & Yuen, 2019; Grenfell *et al*, 2016; Chan & Wong, 2012; Melters *et al*, 2019 in press), and for chaperone interactions required for correct targeting of CENP-A, which we, and others, have demonstrated is defective in cancer cells (Athwal *et al*, 2015; Zhao *et al*, 2016; Nye *et al*, 2018).

## Material and Methods

### Key Resources Table

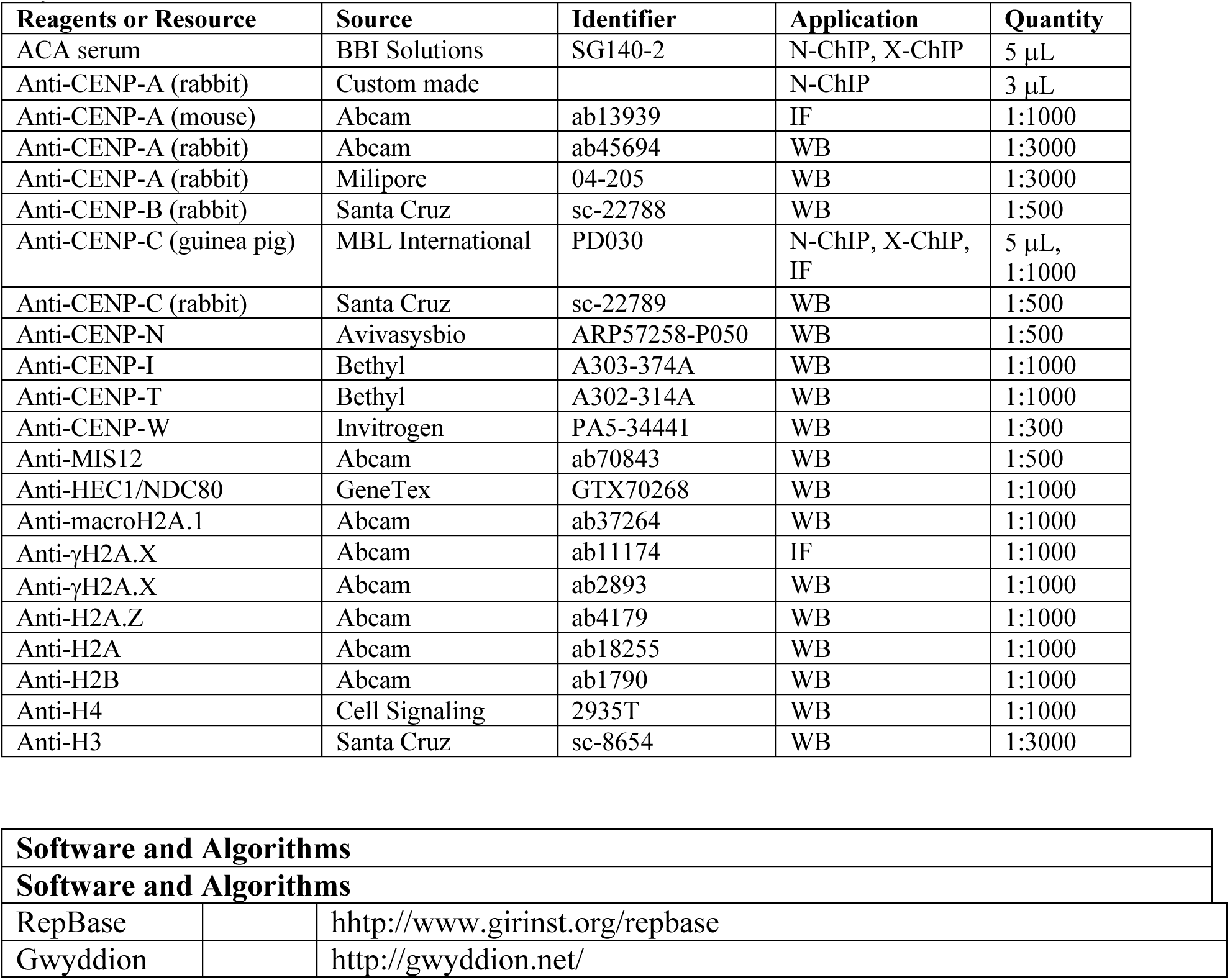
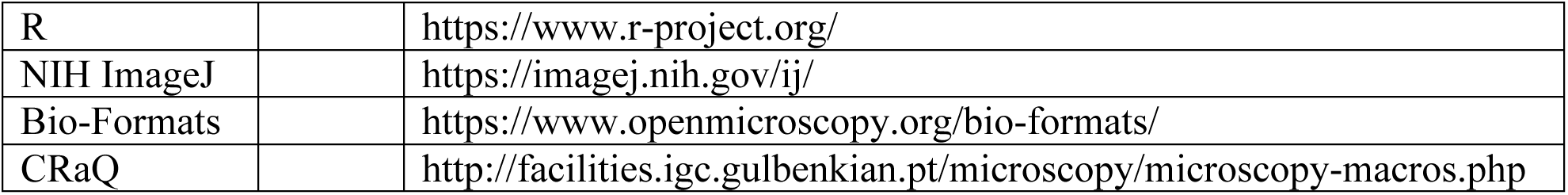

### Native and cross-linked Chromatin-Immunoprecipitation and Western blotting

Human cell line HeLa were grown in DMEM (Invitrogen/ThermoFisher Cat #11965) supplemented with 10% FBS and 1X penicillin and streptomycin cocktail. N-ChIP experiments were performed without fixation. After cells were grown to ∼80% confluency, they were harvested as described (Bui *et al*, 2012, 2017). For best results for chromatin preparation for AFM the pellet that is obtained after each spin-down during the nuclei extraction protocol (Walkiewicz *et al*, 2014a) is broken up with a single gentle tap. Nuclei were digested for 6 minutes with 0.25 U MNase/mL (Sigma-Aldrich cat #N3755-500UN) and supplemented with 1.5 mM CaCl_2_. Following quenching (10 mM EGTA), nuclei pellets were spun down, and chromatin was extracted gently, overnight in an end-over-end rotator, in low salt solution (0.5X PBS; 0.1 mM EGTA; protease inhibitor cocktail (Roche cat #05056489001). N-ChIP chromatin bound to Protein G Sepharose beads (GE Healthcare cat #17-0618-02) were gently washed twice with ice cold 0.5X PBS and spun down for 1 minute at 4°C at 800 rpm. Following the first N-ChIP, the unbound fraction was used for the sequential N-ChIP. X-ChIP experiments were performed with fixation (Skene & Henikoff, 2015). Westerns analyses were done using LiCor’s Odyssey CLx scanner and Image Studio v2.0.

### Glycerol gradient sedimentation

A total of 2 mL of extracted chromatin was applied to 10 mL of 5 to 20% glycerol gradient containing 50 mM Tris-HCl pH 8.0, 2 mM EDTA, 0.1% NP-40, 2 mM DTT, 0.15 M NaCl, and 1X protease inhibitor cocktail layered over 0.4 mL of 50% glycerol. The chromatin was centrifuged with a SW41Ti rotor (Beckman) at 22,000 rpm for 15.5 hours at 4°C. 1 mL aliquots were fractioned from the top, and DNA and protein samples were separated by either 1.2% agarose gel electrophoreses or 4-20% SDS-PAGE gels, respectively. Serial N-ChIP was performed on all 12 fractions.

### AFM and image analysis

Imaging of CENP-C and CENP-A N-ChIP and bulk chromatin was performed as described (Dimitriadis *et al*, 2010; Walkiewicz *et al*, 2014a) with the following modifications. Imaging was performed by using standard AFM equipment (Oxford Instruments, Asylum Research’s Cypher S AFM, Santa Barbara, CA) with silicon cantilevers (OTESPA or OTESPA-R3 with nominal resonances of ∼300 kHz, stiffness of ∼42 N/m, and tip radii of 3–7 nm) in noncontact tapping mode. 10 µl of bulk, CENP-A, or CENP-C chromatin sample was deposited on APS-treated mica (Dimitriadis *et al*, 2010; Walkiewicz *et al*, 2014a). The samples were incubated for 10 min, rinsed gently to remove salts, and dried mildly under vacuum before imaging. Automated image analysis was performed as described in (Walkiewicz *et al*, 2014a) with the only modifications that R software was used instead of Microsoft Excel. A total of six biological replicates were performed for CENP-C experiments and three biological replicates for both the CENP-A and bulk chromatin experiments. Bulk chromatin from the same preparation was imaged in parallel to get the baseline octameric range. For all samples, manual spot analyses were performed to confirm accuracy of automated analyses.

### Immuno-AFM

*In vitro* reconstitution of CENP-A (CENP-A/H4 cat#16-010 and H2A/H2B cat#15-0311, EpiCypher, Research Triangle Park, NC) and H3 (H3/H4 cat#16-0008 and H2A/H2B cat#15-0311, EpiCypher Research Triangle Park, NC) nucleosomes were performed as previously described (Dimitriadis *et al*, 2010; Walkiewicz *et al*, 2014a). Chromatin from HeLa cells were obtained from fractions 6 and 7 of a glycerol density gradient (containing on average tri-, tetra-, and penta-nucleosome arrays). These samples were subjected to immuno-AFM as described previously (M. E. Browning-Kelley *et al*, 1997; Cheung & Walker, 2008; Banerjee *et al*, 2012). An aliquot of each sample was imaged by AFM in non-contact tapping mode. The remainder of the samples were incubated overnight at 4°C with anti-CENP-A antibody (Abcam cat #ab13939) in an end-over-end rotator before being imaged by AFM. Finally, these samples were incubated with anti-mouse secondary antibody (Li-Cor’s IRDye 800CW Donkey anti-mouse IgG cat#925-32212) for an hour at room-temperature in an end-over-end rotator and imaged by AFM in non-contact tapping mode. We analyzed the height profiles of the nucleosomes and antibody complexes as described above.

### Transmission electron microscopy

For transmission electron microscopy (TEM), the N-ChIP samples were fixed by adding 0.1% glutaraldehyde at 4°C for 5 hours, followed by 12-hour dialysis against HNE buffer (10 mM HEPES pH=7.0, 5 mM NaCl, 0.1 mM EDTA) in 20,000 MWCO membranes dialysis cassettes (Slide-A-Lyzer Dialysis Cassette, ThermoFisher cat #66005) at 4°C. The dialyzed samples were diluted to about 1 μg/mL concentration with 67.5 mM NaCl, applied to carbon-coated and glow-discharged EM grids (T1000-Cu, Electron Microscopy Sciences), and stained with 0.04% uranyl acetate. Dark-field EM imaging was conducted at 120 kV using JEM-1000 electron microscope (JEOL USA, Peabody, MA) with SC1000 ORIUS 11 megapixel CCD camera (Gatan, Inc. Warrendale, PA).

### Immunostaining of mitotic chromosomes

HeLa cells were synchronized to mitosis with double thymidine block. Primary antibodies γH2A.X, CENP-C, and CENP-A were used at dilution 1:1000. Alexa secondary (488, 568, 647) were used at dilution of 1:1000. Images were obtained using DeltaVision RT system fitted with a CoolSnap charged-coupled device camera and mounted on an Olympus IX70. Deconvolved IF images were processed using ImageJ. Mitotic defects (lagging chromosomes and/or multipolar spindles) were counted for 83 and 76 cells (mock, GFP-CENP-C, respectively).

### Quench pulse-chase immunofluorescence

To quantify *de novo* assembled CENP-A particles, we transfected HeLa cells with SNAP-tagged CENP-A (generous gift from Dan Foltz) in combination with either empty vector or GFP-CENP-C using the Amaxa Nucleofector kit R (Lonza Bioscience, Walkersville, MD) per instructions. The quench pulse-chase experiment was performed according to Bodor et al^17^. In short, following transfection, cells were synchronized with double thymidine block. At the first release TMR-block (S9106S, New England Biolabs, Ipswich, MA) was added per manufactures instruction and incubated for 30 min at 37°C, followed by three washes with cell culture media. At the second release TMR-Star (S9105S, New England Biolabs, Ipswich, MA) was added per manufactures instructions and incubated for 15 min at 37°C, followed by three washes with cell culture media. Fourteen hours after adding TMR-Star, cells were fixed with 1% paraformaldehyde in PEM (80 mM K-PIPES pH 6.8, 5 mM EGTA pH 7.0, 2 mM MgCl_2_) for 10 min at RT. Next, cells were washed the cells three times with ice cold PEM. To extract soluble proteins, cells were incubated with 0.5% Triton-X in CSK (10 mM K-PIPES pH 6.8, 100 mM NaCl, 300 mM sucrose, 3 mM MgCl_2_, 1 mM EGTA) for 5 min at 4°C. The cells were rinsed with PEM and fixed for a second time with 4% PFA in PEM for 20 min at 4°C. Next, the cells were washed three times with PEM. Cells were permeabilized with 0.5% Triton-X in PEM for 5 min at RT and subsequently washes three times with PEM. Next, the cells were incubated in blocking solution (1X PBS, 3% BSA, 5% normal goat serum) for 1 hr at 4°C. CENP-A antibody (ab13979 1:1000) was added for 1 hr at 4°C, followed by three washes with 1X PBS-T. Anti-mouse secondary (Alexa-488 1:1000) was added for 1hr at 4°C, followed by three 1X PBS-T and two 1X PBS washes. Following air-drying, cells were mounted with Vectashield with DAPI (H-1200, Vector Laboratories, Burlingame, CA) and the coverslips were sealed with nail polish. Images were collected using a DeltaVision RT system fitted with a CoolSnap charged-coupled device camera and mounted on an Olympus IX70. Deconvolved IF images were processed using ImageJ. From up to 22 nuclei, colocalizing CENP-A and TMR-Star foci signal were collected, as well directly neighboring regions. Background signal intensity was deducted from corresponding CENP-A and TMR-Star signal intensity before the ratio CENP-A/TMR-Star was determined. Graphs were prepared using the ggplot2 package for R.

### ChIP-seq

CENP-C N-ChIP followed by ACA N-ChIP was conducted, as well as an IgG N-ChIP and input control as described above. Next, DNA was isolated by first proteinase K treated the samples, followed by DNA extraction by phenol chloroform. The samples were used to prepare libraries for PacBio single-molecule sequencing as described in manufacturer’s protocol (PacBio, Menlo Park, CA). Libraries were produced and loaded on ZWM chip either by diffusion or following size selection of the inserts (> 1000 bp) for all four samples. Subsequently, the reads were sequenced on the PacBio RS II operated by Advanced Technology Center, NCI (Frederick, MD). Sequence reads were mapped to either sequences in RepBase, the consensus sequence used by (Hasson *et al*, 2013), and the consensus sequences used by (Henikoff *et al*, 2015). The sequence data can be found under GEO accession number GSE129351.

### Quantification and statistical analyses

Significant differences for Western blot quantification and nucleosome height measurements from AFM analyses were performed using either paired or two-sided t-test as described in the figure legends. Significance was determined at p<0.05.

## Footnotes

Authors contributions: D.P.M., T.R., and Y.D. designed research; D.P.M., T.R, B.M., S.A.G., and D.S. performed research; D.P.M., T.R., B.M., S.A.G., D.S., and Y.D. analyzed data; D.P.M. prepared figures; and D.P.M. and Y.D. wrote the paper; all authors editing the paper.

The authors declare no conflict of interest.

## Acknowledgements

We thank Drs. Tom Misteli and Sam John, and members of the CSEM laboratory for critical comments and suggestions. We thank Dr. Kerry Bloom for encouraging us to purify kinetochore bound centromeric chromatin. We acknowledge Dr. Andrea Musacchio’s suggestion to test for CENP-C depletion in a previous version of this manuscript.. This work utilized the computational resources of the NIH HPC Biowulf cluster (http://hpc.nih.gov). Y.D. and T.R, B.M., D.S., and D.P.M. were supported by the Intramural Research Program of the Center for Cancer Research at the National Cancer Institute/NIH. S.A.G. was supported by NSF grant 1516999.

